# *homie* and *nhomie* insulators compete with each other for interaction with distant copies, affecting enhancer-promoter interactions and *eve* function

**DOI:** 10.1101/2025.10.28.684994

**Authors:** Miki Fujioka, Wenfan Ke, Paul Schedl, James B. Jaynes

## Abstract

Chromatin insulators, a.k.a. boundaries, separate regions of the chromosome with distinct chromatin characteristics, including distinct histone modifications. This activity affects gene expression by allowing chromatin domains to be stably regulated and maintained. They also block enhancer-promoter interactions and, somewhat paradoxically, can facilitate other enhancer-promoter interactions, particularly when they stitch together distant regions of the chromosome by pairing with specific insulator partners. Here we explore how long-range interactions facilitated by insulator pairing are affected by the presence of two specifically interacting partners. Our results show that when two partners are present, they compete, reducing each other’s effects, suggesting that interactions tend to be limited to two interacting partners at any given time. In particular, when a distant transgenic copy of an *eve* insulator (*homie* or *nhomie*) is present, it can interact with either endogenous insulator. But when one endogenous insulator is removed, the remaining one interacts more strongly with the transgenic copy, biasing the induced enhancer-promoter interactions toward those nearest the remaining endogenous insulator. We also show that removing one or both endogenous *eve* insulators significantly reduces endogenous *eve* function at a critical early stage of development, and that the *eve* Polycomb domain expands in both directions when its insulator boundaries are removed.

## INTRODUCTION

Insulators are one of several elements that influence chromosome architecture. These include enhancers and promoters, which interact with each other, sometimes specifically, by looping out the intervening DNA [1–4]. They also include Polycomb response elements (PREs), which seem to interact with each other relatively non-specifically [5–9]. Another is chromatin modifiers, which include both covalent histone modifiers that can be recruited to specific sequences, resulting in the coalescence of chromatin domains by copolymer cosegregation [10, 11], and ATP-dependent chromatin remodelers that can move nucleosomes around, evict them from the DNA, or help to assemble them into higher order structures that inhibit accessibility to the protein complexes required for gene expression [12].

At least in part by affecting chromosome architecture, insulators do 3 major things to influence gene expression. These are enhancer blocking, facilitation of enhancer-promoter interactions, and barrier activity which blocks the spreading of repressive chromatin along the chromosome. The first two of these seem qualitatively different, and we recently showed that the endogenous *eve* insulator *homie* has multiple sub-domains that to a limited extent differentially influence these two activities [13].

In principle, insulators could interact with each other in more than one way, with different consequences for gene expression. First, because there are two copies of each DNA sequence near each other during G2, on the paired sister chromatids, these would be expected to pair with each other, perhaps holding sister chromatids together in register until chromosome segregation during mitosis. Furthermore, when homologous chromosomes are paired, as they are throughout most of the Drosophila life cycle, there are 4 copies of each insulator near each other in G2. We know that paired homologs undergo transvection [14, 15], in which enhancers on one homolog affect gene expression on the other, and that insulators that specifically pair with each other can mediate this effect [16, 17]. This provides the opportunity for multiple copies of a “single” insulator to interact with each other, at the same time that they interact with other, nearby or distant, insulators. On the other hand, having more than one potential partner may result in a “time-sharing” competition between alternative partners.

Consistent with the possibility of an insulator interacting with multiple potential partners, insulators can collect together in insulator bodies [18, 19], which could involve higher order “group” interactions among insulators. When and if more than two insulators interact with each other at the same time, these interactions could be cooperative. Alternatively, only pairwise interactions may occur at any given time, so that multiple potential partners may compete with each other. Indications from artificial insulator bypass and insulator competition studies are more suggestive of competition than of cooperativity when multiple partners are present nearby each other in *cis* [20–22]. Such competition is not inconsistent with insulator body formation, for two reasons. First, collecting a group of insulators together would make it quicker for each of them to find another partner when they dissociate, and this could provide an effectively “attractive” force reducing the average distance between them over time. Second, dissociation of an existing pair might be facilitated by an incipient interaction with a third potential partner, resulting in a “transition state” with all 3 transiently engaged. Such interactions would be facilitated within an insulator body, again providing a force tending to keep the insulator body together.

Here, we use the specifically pairing insulators that flank the *eve* locus to test the effects of providing additional potential pairing partners in a native context (Fig. 1A). We previously established that the endogenous *eve* insulators interact with each other in a specific orientation (head-to-tail) [23], and that they also interact with transgenic copies of themselves and each other from considerable distances away along the chromosome [16]. We also showed that transgenic copies can interact other transgenic copies from great distances in cis (∼3 Mb) [24], as well as in trans, on paired homologous chromosomes (∼2Mb) [16]. These long-range chromosomal interactions facilitate enhancer-promoter (E-P) interactions in a manner that depends on whether the elements are on the same side of the paired insulators or not, and on how close the enhancer-promoter pairs are to the interacting elements. While these studies using endogenous *eve* insulators and enhancers in combination with transgenic copies of either *homie* or *nhomie* adjacent to reporter gene promoters taught us a great deal about how E-P facilitation and blocking are affected by insulator-induced chromosome topologies [13, 16, 23, 25, 26], they were complicated by the fact that there were two partners for transgenic insulator copies, namely endogenous *homie* and *nhomie*. To simplify the system, we removed one endogenous *eve* insulator at a time, and investigated how this changes the resulting E-P interactions. Our new results are suggestive of competition, indicating that pairwise interactions may be predominant, at least for this type of insulator, where pairing is highly specific (and directional), with either an endogenous partner, or with copies of themselves.

**Figure 1.**
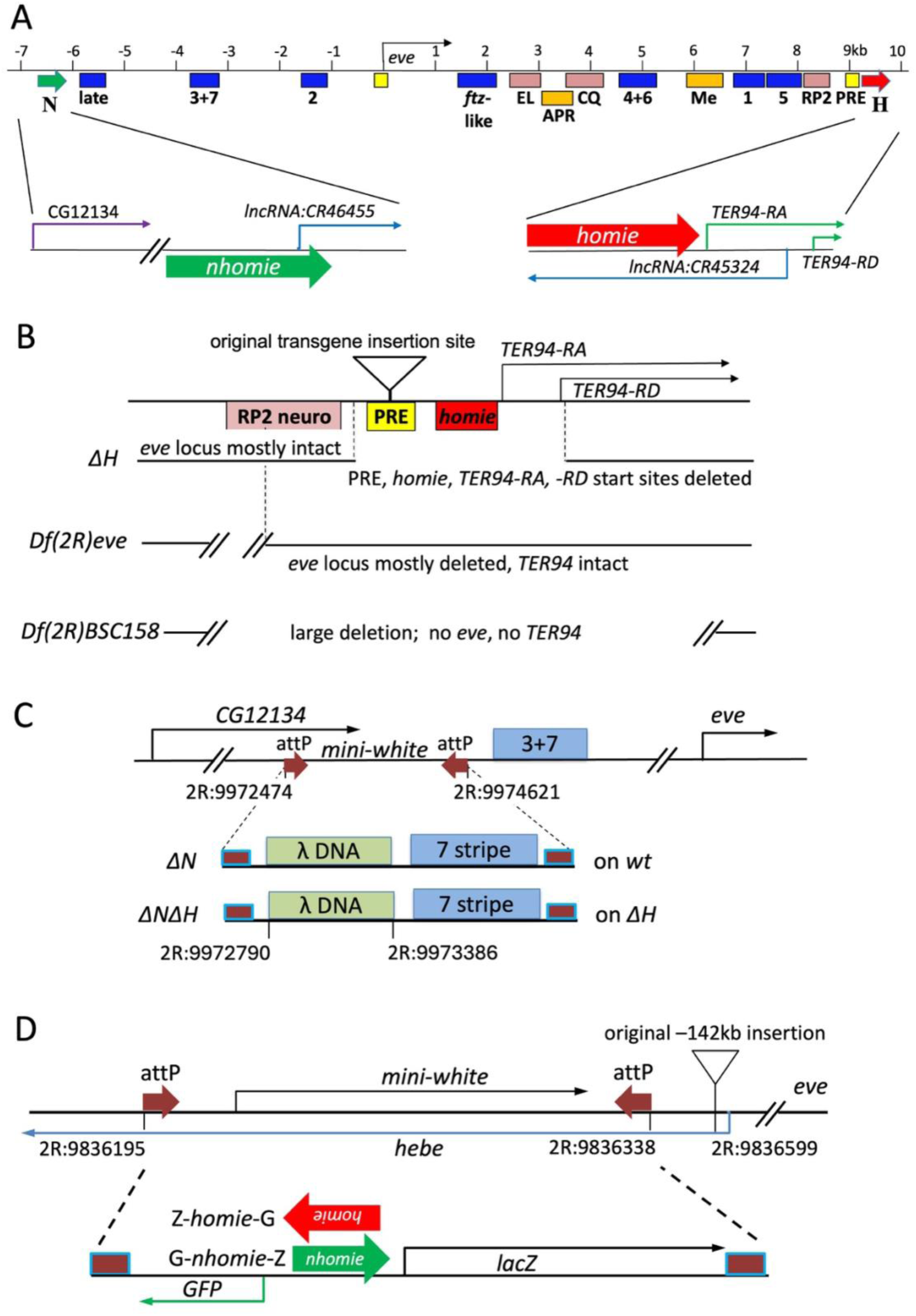
Maps of modified chromosomes and transgenes used in this study. **A.** Map of the *eve* locus. Boxes indicate locations of enhancers. Blue: stripe enhancers (early stripes: 3+7, 2, 4+6, 1, 5; 7 late stripes: late; *ftz*-like stripes: *ftz*-like) [28–33], pink: neuronal enhancers (EL, CQ, and “RP2”, which drives expression in RP2+a/pCC cells) [28, 29], orange: anal plate ring (APR) and mesodermal (Me) enhancers [28, 29], yellow: PREs [61]. Green block arrows: *nhomie* [16]. Red block arrows: *homie* [16, 24]. Start sites and directions of transcripts are shown as thin arrows. **B.** Maps of mutant chromosomes used in this study. *ΔH*: small deletion (2R: 9987987.. 9989353 is deleted) caused by imprecise excision of a homed P-element transgene (insertion site: triangle). It removes the *eve* 3’ PRE, *homie*, and the transcription start sites of the *TER94-RA* and -*RD* transcripts. *Df(2R)eve* [27, 36, 54]: the deletion’s 5’ end is in *Mef2*, and its 3’ end is in the middle of the RP2 neuronal element (2R: 9949630.. 9987407 is deleted, see sequence in Fig. S1). Most of the *eve* locus is deleted, but *TER94* is intact. *Df(2R)BSC158*: large deletion (2R:9875312..10025310 is deleted) [34, 35]. The *eve* and *TER94* loci are removed. *eve^R13^* [36]: an amorphic point mutation in the *eve* coding region, which causes premature termination of the Eve protein [29]. **C.** Map of *ΔN*, the *nhomie* deletion. The top line shows the attP insertion created using CRISPR. The *mini-white* gene was then replaced by the DNA fragment shown below, using RMCE. The 600bp *nhomie* region was replaced by a similar-sized fragment of λ DNA (2R: 9972790..9973386 is deleted) on either a wild-type (creating *ΔN*) [23] or the *ΔH* chromosome (creating *ΔNΔH*). The end points used to create the deletion are shown on the map. Brown square with blue outline: attP/attB fusion sequence left after RMCE [58]. **D.** Maps of reporter transgenes. Top map shows the attP-carrying chromosome used for RMCE. The original insertion site at –142kb [16, 24] is shown as a triangle (2R:9836599). Dual reporter genes carrying either *homie* (Z-*homie*-G) or *nhomie* (G-*nhomie*-Z) replaced *mini-white*. *homie* and *nhomie* in the transgene are oriented so that *lacZ* is more strongly expressed when they interact orientation-specifically with the *eve* locus (see text).

## RESULTS

### Long-range insulator interactions can facilitate locally biased enhancer-promoter interactions

Previous studies showed that *homie* and *nhomie* can mediate long-distance regulatory interactions when included in a reporter transgene that is inserted within several Mb of the *eve* locus [16, 24]. Because the pairing interactions of the *eve* insulators are orientation-dependent, one of the two reporters in a dual reporter transgene is preferentially activated by the *eve* enhancers [16]. Although there are *eve* enhancers located both upstream (on the *nhomie* side) and downstream (on the *homie* side) of the *eve* transcribed region (Fig. 1A), there is no apparent bias in their ability to activate reporter expression. This is reflected in the pattern of physical contacts seen in MicroC experiments, which extend across the entire *eve* TAD [25]. Moreover, when the transgene insulator is used as a viewpoint, it contacts both *nhomie* and *homie* [25]. One interpretation of the MicroC data is that the transgene insulator interacts simultaneously with the two endogenous *eve* insulators, forming a tripartite complex. An alternative possibility is that there are three different pairwise interactions, transgene insulator with either endogenous *nhomie* or *homie*, and endogenous *nhomie* with *homie*, generating three different types of loops that potentially compete with each other over a fairly short time period (on the order of tens of minutes or less). This made us wonder how the transgene would interact with the *eve* enhancers if there were only one endogenous *eve* insulator.

### Endogenous *homie* and *nhomie* are required for the long-range interaction

To answer this question, we first deleted either endogenous *nhomie* or *homie*. We used CRISPR to remove endogenous *nhomie* (creating *ΔN*; see Fig. 1C for map)[23], but were unsuccessful at obtaining a similarly “clean” removal of endogenous *homie*. However, we were able to obtain a small deletion that includes endogenous *homie*, along with the 3’ *eve* PRE and the *TER94* basal promoter region and first exon (Fig. 1B; see Fig. S1 for sequence). This mutant chromosome (*ΔH*) is homozygous lethal after embryogenesis. In order to test whether the lethality of *ΔH* comes from removal of *homie* or the *TER94* promoter, we complemented it with an *eve*-deficient chromosome carrying an intact *TER94* locus. To do this, we first confirmed that *Df(2R)eve* is a complete null for *eve* function [27], but has intact *TER94* function (Fig. 1B; see Fig. S1 and Materials and Methods for details). When the *ΔH* chromosome was placed over *Df(2R)eve*, some adult flies survived. Some of the surviving adults showed abnormal segmentation that is not evident in *Df(2R)eve* heterozygotes. Nonetheless, rescue to adulthood by *Df(2R)eve* indicates that the recessive lethality of *ΔH* is mainly caused by lack of *TER94* function. Segmentation defects in *ΔH/Df(2R)eve* adult flies suggest that the *eve* function provided by *ΔH* is somewhat weakened, and this is confirmed below by analyzing embryonic defects. For further analysis, we generated a chromosome lacking both *homie* and *nhomie* (*ΔNΔH*), replacing *nhomie* on the *ΔH* chromosome with *lambda* DNA, using CRISPR (Fig. 1C; see also Materials and Methods). The fertility of *ΔNΔH* seems to be lower than that of *ΔN* or *ΔH*, as we could maintain the *ΔNΔH* line for only a few generations.

In order to analyze interactions between transgenes and the modified endogenous *eve* locus, we used CRISPR to insert a pair of attP sites (flanking a *mini-white* marker gene) onto the wild-type (wt), *ΔN*, *ΔH,* and *ΔNΔH* chromosomes, near our previously used attP site (142kb upstream of *eve*). The *mini-white* marker was then replaced with our *lacZ-GFP* dual reporter transgene vector [13, 16] (Fig. 1D, see Materials and Methods). In previous studies [16, 25], we described how transgene activation was biased according to how *homie* or *nhomie* was oriented in the transgene relative to the two reporter genes. The reporter located either upstream of *homie* or downstream of *nhomie* is preferentially activated (see Fig. 1A, D; red or green arrow shows the orientation of *homie* or *nhomie*, respectively). Throughout this study, we used the transgenes that preferentially express *lacZ*, which we call Z-*homie*-G and G-*nhomie*-Z. The negative control (*lambda* DNA in place of a transgenic insulator) did not show any *eve*-like *lacZ* expression (Fig. S2). However, various patterns of ectopic expression that we previously reported [13] were seen. Fig. S3 shows positive control expression of each of the two reporters with *homie* and *nhomie* at the new attP site on a wt chromosome. In both lines, *lacZ* is expressed as 7 stripes at stages 5-9 (Fig. S3, stages 5 and 7 are shown), and in a tissue-specific *eve* pattern in the mesoderm, anal plate ring (APR), and central nervous system (CNS) at later stages (Fig. S3, stages 11 and 13 are shown). *GFP* is not expressed in the *eve* pattern, instead, ventral midline expression in the CNS driven by an enhancer near the insertion site is observed. In each case, these expression patterns match previously reported results with the same dual-reporter transgenes inserted at the original attP site [13, 16].

In previous experiments, we found that long-range (LR) interactions between reporters in the transgene and the *eve* enhancers only take place when either *homie* or *nhomie* is included in the transgene [13, 16]. We concluded that the LR interaction between the *eve* locus and transgenic reporter genes was mediated by directional (orientation-specific) pairing between *homie*/*nhomie* in transgenes and their endogenous counterparts. Further analysis using live-cell imaging [26] and physical interaction studies using MicroC [25] were consistent with this explanation. This model predicts that the LR interaction will disappear without endogenous *homie* and *nhomie*. So, we tested whether removing both endogenous *homie* and *nhomie* (*ΔNΔH*) causes loss of the LR interaction. As predicted, no *eve*-like *lacZ* expression was seen from either Z-*homie*-G or G-*nhomie*-Z when they were carried on *ΔNΔH* (Fig. 2, compare wt and *ΔNΔH*). These findings indicate that physical pairing between the endogenous and transgenic copies of the *eve* insulators are required for the LR interaction.

**Figure 2.**
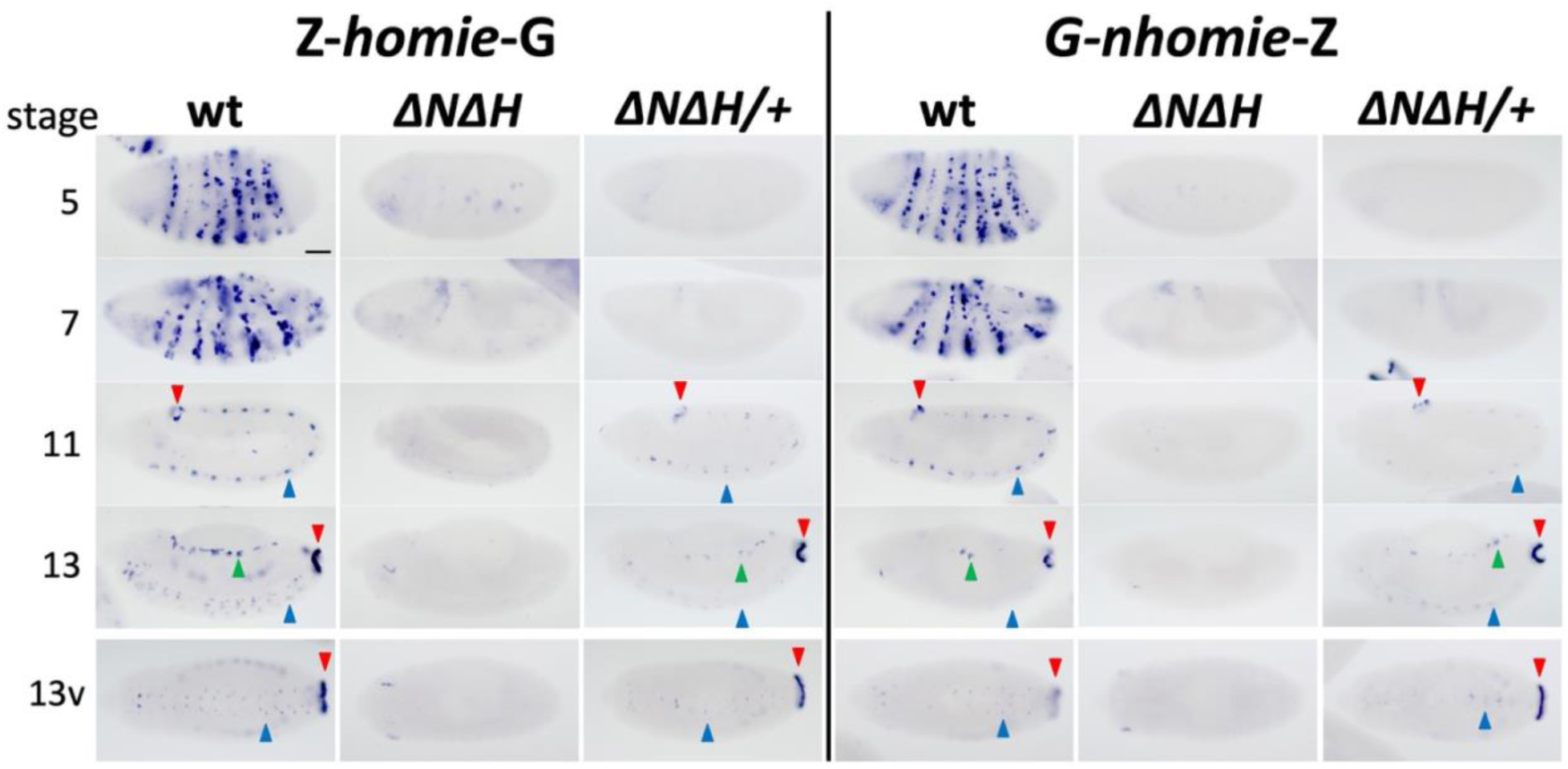
Transgenic *homie* and *nhomie* require endogenous *homie* and *nhomie* for long-range interaction with *eve* enhancers, which can occur in trans at later embryonic stages. RNA expression from *lacZ* reporter transgenes carrying *homie* (*Z-homie-G*) or *nhomie* (*G-nhomie-Z*) located at –142kb on homozygous wild-type (wt) and *ΔNΔH* chromosomes, and heterozygous *ΔNΔH* over wild-type (ΔNΔH/+, showing a *trans* interaction). Arrowheads indicate *eve* tissue-specific expression: red: APR; blue: CNS; green: mesoderm. Embryonic stages 5, 7, 11, and 13 are shown (ventral views in bottom row). Scale Bar: 50μm.

To test this idea further we took advantage of the fact that once homologs in somatic cells pair with each other, trans-regulatory interactions (transvection) are observed if enhancers on one homolog are brought into close proximity with target genes on the other homolog. Thus, if insulators at the *eve* locus are required for interactions with the transgene, then it should be possible to restore *lacZ* expression with a wild-type *eve* locus is *trans* to either Z-*homie*-G,*ΔNΔH* or G-*nhomie*-Z,*ΔNΔH.* Homolog pairing is limited at the blastoderm stage, and as expected we do not detect *eve*-like *lacZ* expression. However, later in development, during stages 11-13, APR enhancers drive *lacZ* expression in the posterior, and expression in the CNS and mesoderm are also detected (Fig. 2, compare *ΔNΔH* and *ΔNΔH/+*).

### Pairwise insulator interactions bias enhancer-promoter interactions

We next asked whether endogenous *homie* and *nhomie* individually can mediate the LR interaction with transgenic copies of either themselves or each other. To do this, *mini-white*-carrying attP sites were introduced on each of the *ΔH* and *ΔN* chromosomes as described above (Fig. 1D). On either *ΔN* or *ΔH*, Z-*homie*-G and G-*nhomie*-Z each showed an LR interaction, expressing *lacZ* in a partial *eve* pattern. Importantly, however, these patterns differed from the interactions seen with the intact *eve* locus (Fig. 3). In wild-type embryos, at stage 5, each stripe is controlled by a distinct enhancer [28–33], while 7 stripes at stages 6-8 are under control of a different “7-stripe” or “late stripe” enhancer [32, 33] (see Fig. 1A for enhancer locations). In a wt background, at stage 5, the level of *lacZ* expression from either Z-*homie*-G or G-*nhomie*-Z is close to equal among the different stripes. (We note that the stripe 2 enhancer typically drives less *lacZ* expression than the others at this stage, which could be due to promoter competition, as the stripe 2 enhancer is located next to the endogenous *eve* promoter.) At stage 7, the 7 stripes are also expressed at similar levels.

**Figure 3.**
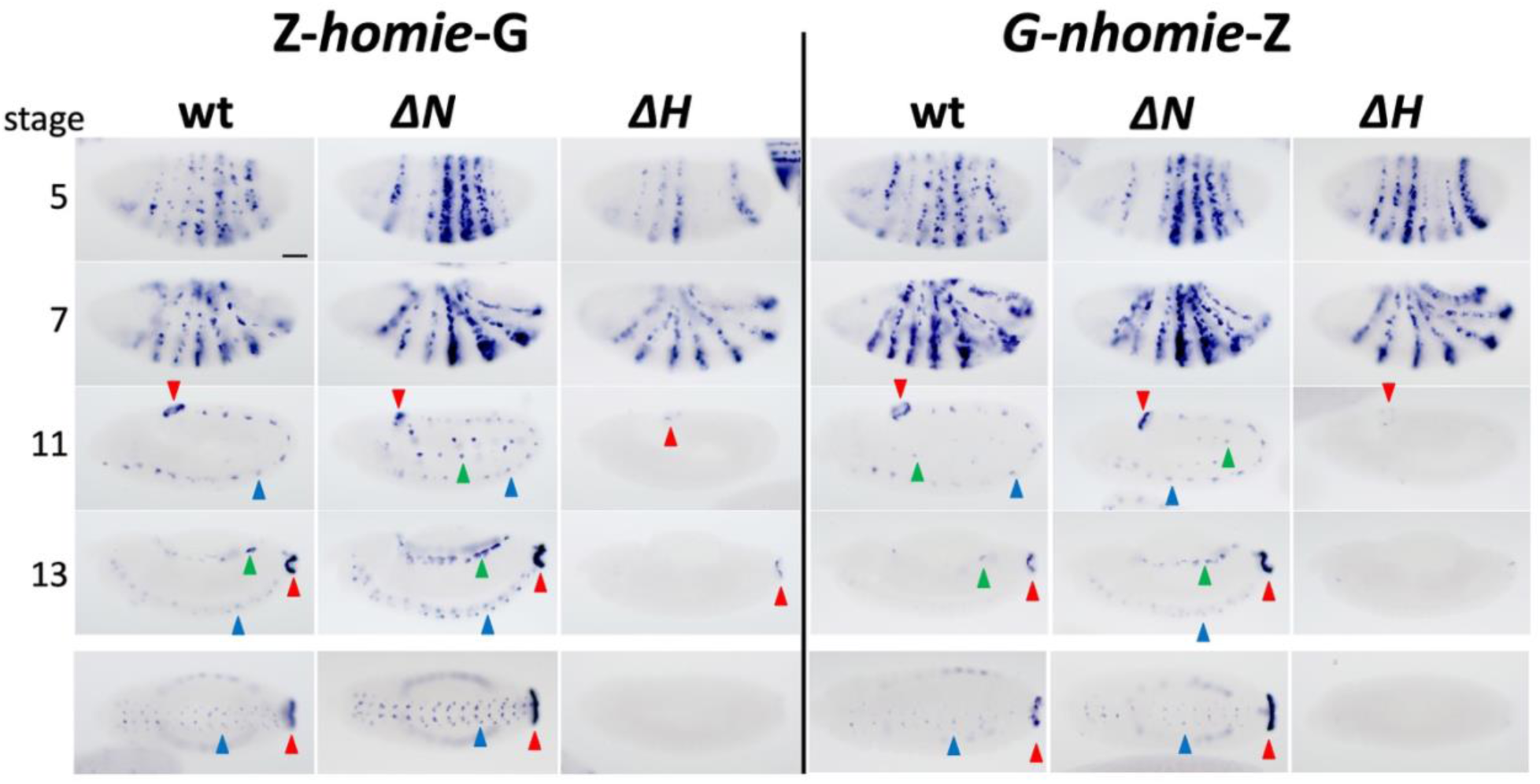
When one endogenous *eve* insulator is removed, long-range interactions become biased toward enhancers near the remaining endogenous insulator. RNA expression from *lacZ* reporter transgenes carrying *homie* (*Z-homie-G*, left panel) or *nhomie* (*G-nhomie-Z*, right panel) located at –142kb on homozygous wild-type (wt), *ΔN*, and *ΔH* chromosomes. Arrowheads indicate *eve* tissue-specific expression: red: APR; blue: CNS; green: mesoderm. Embryonic stages 5, 7, 11, and 13 are shown (ventral views in bottom row). Scale Bar: 50μm.

The pattern of *lacZ* expression in the *ΔN* differs from that in wt in that the level of *lacZ* expression in the seven stripes at stage 5 is much more uneven. With either Z-*homie*-G or G-*nhomie*-Z on *ΔN*, *lacZ* is expressed strongly in stripes 4, 5, and 6 and more weakly in stripes 1, 2, 3, and 7 (Fig. 3, *ΔN*). At stage 7, the 7-stripe pattern is observed, but stripes 4, 5, and 6 appear stronger because there is expression persisting from stage 5. The *eve* tissue-specific pattern in mesodermal, CNS, and APR are also observed, and the level of *lacZ* expression appears stronger than with the intact *eve* locus (Fig. 3, compare wt and *ΔN*). Strikingly, the enhancers for each of these aspects of the *eve* pattern are located 3’ of the *eve* transcription unit, closer to the remaining endogenous insulator, *homie* (Fig. 1A).

In contrast, with *ΔH* at stage 5, *lacZ* expression from either Z-*homie*-G or G-*nhomie*-Z is stronger in stripes 1, 2, 3, and 7, and weaker in stripes 4, 5, and 6. This is reciprocal to the pattern with *ΔN*. The 7-stripe expression at stage 7 is similar to that seen in wild type. Furthermore, the later-stage tissue-specific expression at stages 11-13 is less frequently observed, and then only faintly (Fig. 3, compare wt and *ΔH*). The enhancers for stripes 2, 3, and 7, as well as the 7-stripe enhancer, are all located upstream of the *eve* coding region, closer to *nhomie* (Fig. 1A). We also note that stripe 1 enhancers reside both upstream [33] and downstream of the coding region [28, 29].

A straightforward possibility suggested by these results is that *homie-* or *nhomie*-carrying transgenes engage in LR interactions with either endogenous *homie* alone or *nhomie* individually, rather than at the same time. When they do so, their reporters are activated more strongly by the *eve* enhancers that are closer (on the same side of the *eve* transcription unit) than when the endogenous locus carries both *homie* and *nhomie*. When the endogenous *eve* locus is intact, the expression of *lacZ* is essentially a composite of the patterns seen in *ΔN* and *ΔH*, but weakened somewhat relative to the sum of these two individual patterns, consistent with some competition between the endogenous insulators for the transgenic copy when both are present.

### Transgene – endogenous *eve* interactions: cis vs. trans

If insulator pairing interactions are predominantly pairwise, then there should be a competition between potential pairing partners when there are three or more of them. To test this idea, we manipulated the number of competing insulators by generating trans combinations between homologs carrying the transgene and different wild type and mutant versions of the *eve* locus. We focused on whether later-stage interactions become stronger without a trans copy of the *eve* locus, which would be consistent with competition occurring. To remove the trans copy of endogenous *eve*, we used the large chromosomal deficiency *Df(2R)BSC158* (*BSC158*) [34, 35]. At stages 5-7, the stripe expression was not clearly increased. However, we did see an increase in expression in the CNS at later stages with the deficiency, when homologs are paired, with either *nhomie* or *homie* in the transgene (Fig. 4: cis interaction only, 2^nd^ and 5^th^ columns, vs. in cis with a trans copy of *eve* on a *Sco*-carrying chromosome, 1^st^ and 4^th^ columns). However, this is probably not due to removal of competing insulators, because when we use the *ΔNΔH* chromosome in trans to remove the potentially competing copies of *eve* insulators, we do not see the effect (Fig. 4, 3^rd^ column). So, the effect is probably not due to “trans competition” between the transgene and the trans copy of the endogenous insulators for the cis copy of the endogenous locus. It is more likely due to the lack of chromosome pairing when the *BSC* deficiency is present in trans (see Discussion).

**Figure 4.**
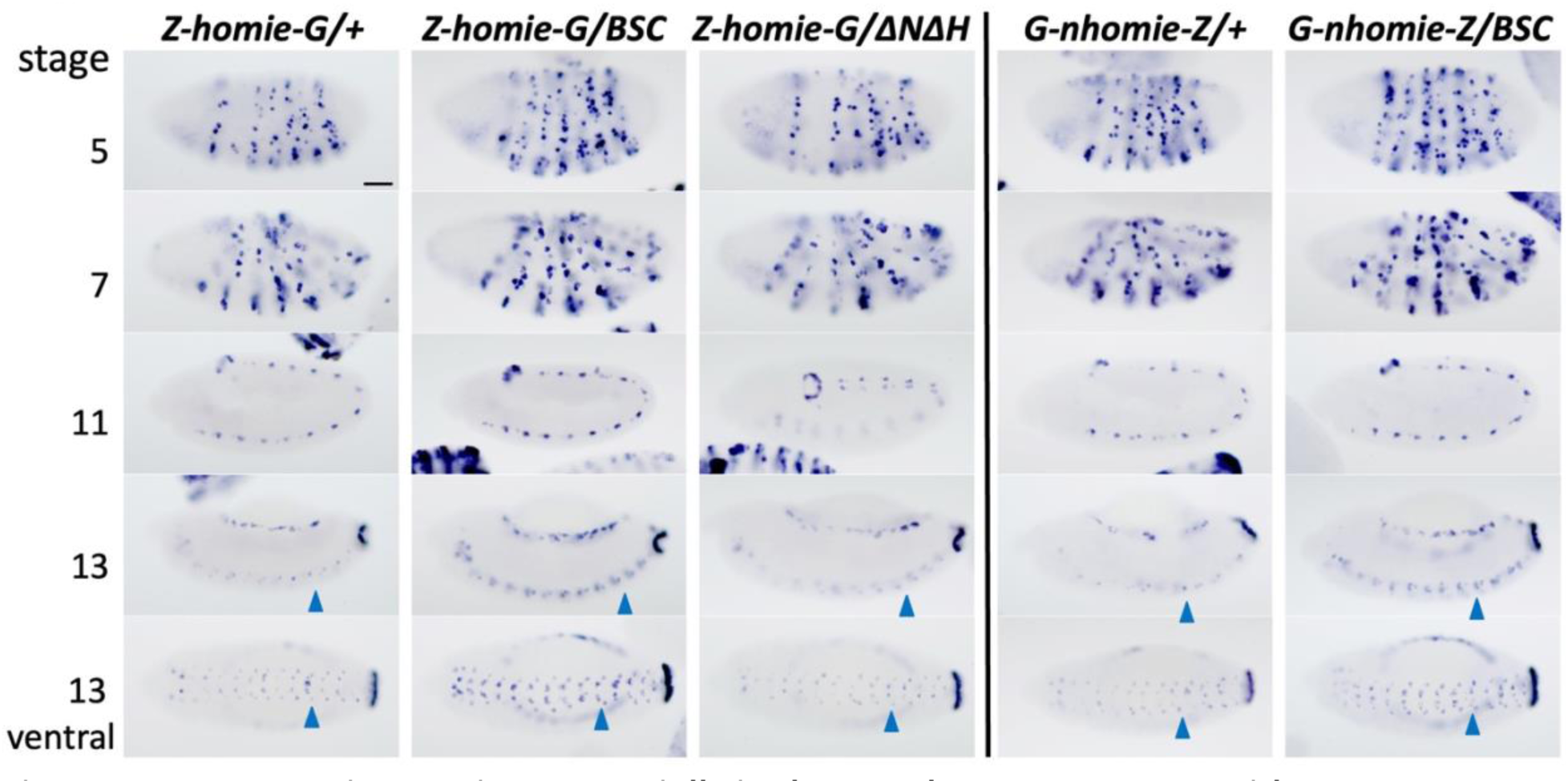
Late-stage interactions, especially in the CNS, become stronger without a trans copy of the *eve* locus. RNA expression from heterozygous *lacZ* reporter transgenes carrying *homie* (*Z-homie-G*, left panel) or *nhomie* (*G-nhomie-Z*, right panel) located at –142kb, in trans with either a chromosome carrying a wild-type *eve* locus (reporter transgene/+), *Df(2R)BSC158* (reporter transgene*/BSC*), or *ΔNΔH* (*Z-homie-G/ΔNΔH*). Embryonic stages 5, 7, 11, 13 are shown (ventral views in bottom row). Note that *lacZ* expression, especially in the CNS at stage 13 (blue arrowheads), is stronger when the *eve* locus is absent in trans (compare *BSC* to *+*), but not when only the *eve* insulators are missing on the trans chromosome *(ΔNΔH*). Scale Bar: 50μm.

### Effects of endogenous insulator removal on *eve* function

In a previous study using the *nhomie* deletion *ΔN*, we found that homozygous mutants were viable and fertile. However, *ΔN* is weakly haploinsufficient, so that when in trans to the *BSC158* deletion, nearly 10% of embryos were missing two or more ventral abdominal denticle bands [23]. To extend this analysis, we have now analyzed embryos derived from self-crossing *ΔNΔH*/*SM6a,Cy*, Δ*H/SM6a,Cy*, and homozygous *ΔN*, for cuticular segmentation defects. As controls, we used *Sco/SM6a,Cy* (“*Sco/Cy*”), controlling for the *SM6a* chromosome in two of the lines, and *yw*, as the control for the *ΔN* self-cross. In all these cases, more than 98% of embryos showed a wild-type cuticle phenotype (Fig. S4). In contrast, embryos from a cross of *Sco/Cy* x *BSC158/Cy* showed a missing ventral abdominal denticle band (most of them in the abdominal segment A6) in ∼10-25% of embryos (Fig. 5, “WT” in “% affected”). The *Sco/BSC158* and *Cy/BSC158* progeny (which each have a single copy of *eve*) together represent about 50% of these embryos, so this phenotype is not fully penetrant. However, the fact that progeny from the above self-crosses did not show such segmentation defects indicates that homozygotes for the insulator deletions have more than one wild-type copy’s worth of normal *eve* activity (see below and Discussion).

**Figure 5.**
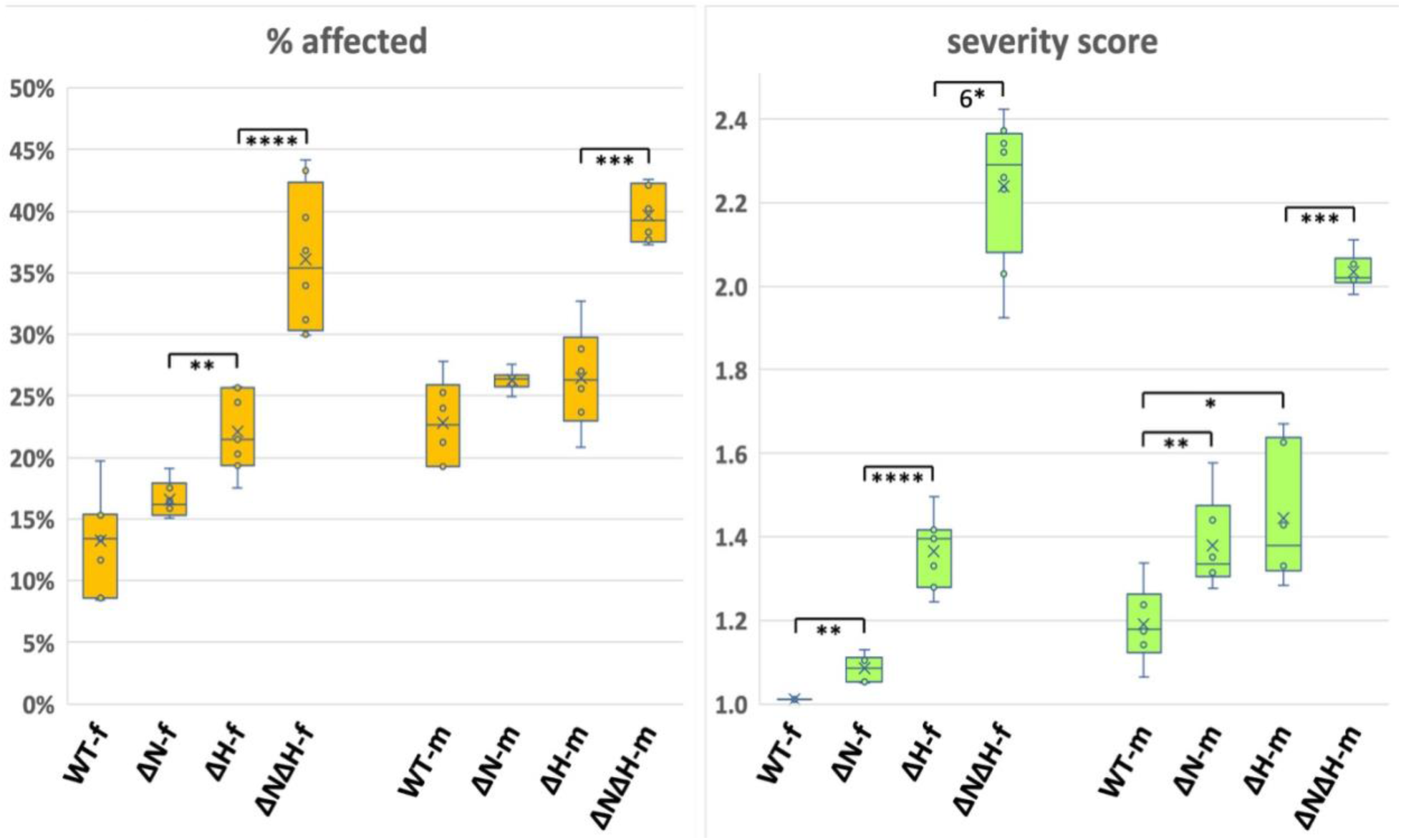
Deleting both *nhomie* and *homie* from endogenous *eve* compromises function more than does deleting only *homie* or *nhomie*, when the genetic background is sensitized to changes in *eve* function. *Sco/Cy* (WT)*, ΔN/Cy* (ΔN)*, ΔH/Cy* (ΔH) and *ΔNΔH/Cy* (ΔNΔH) lines were crossed with *Df(2R)BSC158/Cy*, and embryonic cuticle defects were counted. The direction of the cross is indicated as -f: female and -m: male. The percentage of embryos showing segmentation defects (left graph, % affected) and the average number of ventral abdominal denticle bands deleted per non-wild-type embryo (right graph, severity score) are shown as box-and-whiskers plots. The pair-wise significance of differences (p-values) is indicated as: *: p < .05, **: p < .01, ***: p < .001, ****: p < .0001, 6*: p < 10^−6^. The data are from two separate 3-way comparison experiments, involving [*Sco/Cy, ΔN/Cy,* and *ΔH/Cy*] and [*Sco/Cy, ΔH/Cy*, and *ΔNΔH/Cy*], and these two data sets were combined as described in Materials and Methods. The number of cuticle preparations used (n) and the total number of embryos counted were as follows: WT-f (n=7, 1414 embryos), ΔN-f (n=6, 1676), ΔH-f (n=7, 1885), ΔNΔH-f (n=8, 1391), WT-m (n=6, 1311), ΔN-m (n=6, 1676), ΔH-m (n=6, 1928), ΔNΔH-m (n=6, 2541).

Since we found that *ΔN* is weakly haploinsufficient, we crossed each of the three insulator deletion lines with a series of different *eve* mutant chromosomes, as diagrammed in Fig. 1B. For Δ*nhomie*, we used Δ*N*/*SM6a,Cy* in order to make the crosses equivalent, even though the *ΔN* lines are homozygous viable and fertile. To compare the number and severity of segmentation defects, we calculated two measures: the percentage of affected embryos in each population (Fig. 5, % affected) and the average number of ventral abdominal denticle bands deleted in the affected population (Fig. 5, severity score). When placed over the *BSC158* chromosome, both *ΔH* and *ΔNΔH* increased the percentage of embryos showing segmentation defects significantly over that of the control *Sco/Cy* (“WT”, Fig. 5) in at least one direction of the cross. In order to present all of the data with this set of crosses together, the data from the experimental set [WT, *ΔN,* and *ΔH*] were merged with those from the experimental set [WT, *ΔH*, and *ΔNΔH*] to produce Fig. 5 (see Materials and Methods). The statistical analysis used the WT data from the latter experiment, which had a larger variance than the WT data from the former experiment. In the former experiment, *ΔN* did show a significant increase in percentage of defects over WT (Fig. S5A), even though this same comparison in the merged data set did not show a significant difference (Fig. 5). Importantly, the severity of defects in each of the 3 cases was significantly increased over that of WT. Furthermore, both the percentage of defective embryos and the severity of segmentation defects were greater with *ΔNΔH* (Fig. 5) than with either *ΔN* or *ΔH* alone.

Since the *BSC158* deficiency is large and encompasses many other genes besides *eve*, we further tested these effects for *ΔH* and *ΔNΔH* using both *Df(2R)eve* and *eve^R13^*[36]. *Df(2R)eve* deletes most of the *eve* locus plus several kilobases upstream (see sequences of junction fragments in Fig. S1), while *eve^R13^* is a point mutant causing premature termination of the Eve protein [29]. The effects of combining these two *eve* mutants with the boundary deletions are similar to those observed for the *BSC158* deficiency, indicating that they are due to the absence of one functional copy of *eve* in combination with the loss (in trans) of one or both *eve* insulators in the remaining, now partially functional, copy (Figs. S5B, C, and S6, which uses the same data shown in Figs. 5, S5B, and S5C, but presented as stacked graphs for easy visualization of the differences). We also note that in the *Df(2R)eve* crosses, the effect of *ΔNΔH* (Figs. S5B and S6B) is significantly more severe when it comes from the female parent, for unknown reasons.

In all of the cases described above, virtually all of the missing abdominal denticle bands occurred in even-numbered segments (A2, A4, and A6). This is significant because it is mostly the even-numbered segments that are deleted in *eve* deficiency mutants [27, 36]. This indicates that the defects are most likely due to a loss of *eve* function in early embryos.

### *eve and engrailed* expression are disrupted in *eve* insulator deletion mutants

The segmentation defects in embryos trans-heterozygous for the insulator deletions and the different *eve* mutants are expected to be due to alterations in the pattern of *eve* expression during the blastoderm stage. Fig. S7A shows the pattern of *eve* expression detected by *in situ* hybridization in three representative wt, *+/BSC158*, and *ΔNΔH/ BSC158* stage 5 embryos. As expected from the incomplete penetrance of the segmentation defects in *+/BSC158* embryos, *eve* expression in many *+/BSC158* embryos is indistinguishable from the wt control, except for generally weaker expression. However, in a subset of the *+/BSC158* embryos, expression in stripes 5 and 6 is more severely reduced. The effects on *eve* expression are more pronounced in *ΔNΔH/ BSC158* embryos, in that *eve* expression is more clearly reduced in all of the stripes, and this reduction is typically greater in stripes 4, 5, 6, and 7.

Embryos from these crosses were then analyzed for *engrailed* (*en*) expression, which is a critical downstream target of *eve* (Fig. S7B) [37–40]. We examined the pattern of *en* expression in wt, *+/BSC158*, and *ΔNΔH/BSC158* at different stages of development. In *+/BSC158* embryos, which has only one copy of endogenous *eve*, *en* expression is abnormal from the time it is initiated. Instead of a relatively even spacing of the 14 *en* stripes, the stripes are “twinned” (Fig. S7B, compare wt and +/*BSC158*). This is due to decreased *eve* expression, which causes a narrowing of the parasegments that span the 7 early *eve* stripes, by a mechanism that has been described in detail [41–45]. Numbers in Fig. S7B (stage 7) show the positions of these 7 *eve* stripes relative the 14 *en* stripes. As a downstream consequence of this narrowing, there is occasional “degeneration” of the parasegment at later stages, resulting in ectopic and/or haphazard *en* expression within the narrowed parasegment (arrowheads in stage 13 embryos). This type of defect is the presumed precursor of the cuticle defects quantified in Figs. 5, S5, and S6. An example of an embryo with such defects in 4 adjacent parasegments is shown in Fig. S7B, 3^rd^ column (*ΔNΔH/BSC*, stage 13), presumably representing an embryo that will develop the severe pattern of defects with 4 denticle bands deleted, a phenotype only rarely observed in the *+/BSC* control.

### Endogenous insulator removal allows the *eve* Polycomb domain to spread

In previous studies, we found that the *homie* boundary blocked the spread of the repressive Polycomb histone modification H3K27me3 [13]. We also showed that removing *homie* from a modified transgenic *eve* “pseudo-locus” (in which the *eve* coding region is replaced with a *lacZ* reporter gene, and the *TER94* coding region is replaced with a *GFP* reporter) caused the spreading of H3K27me3 into *TER94*, and reduction of *TER94-GFP* transcript levels [46]. Those data suggested that *homie* prevents the spreading of repressive H3K27me3 in order to prevent the repression of *TER94*. Here, we tested whether we could see similar spreading of H3K27me3 outside of the endogenous *eve* locus in the absence of *nhomie* and *homie*, using our *ΔNΔH* mutants. It is important to note that this experiment differs from the transgenic experiment in two important respects. First, *ΔNΔH* is homozygous in only 1/4 of the embryos from the cross and is heterozygous in another 1/2 of the population, so H3K27me3 is expected to increase outside the *eve* locus in at most 1/2 of the chromosomes being analyzed in the experiment. Second, *ΔNΔH* is also missing the *eve* 3’ PRE, which might cause a reduction in the ability of the *eve* Polycomb domain to spread. Despite these differences, both of which are expected to reduce Polycomb spreading, we were able to see a significant increase in H3K27me3 outside the endogenous *eve* locus in both directions due to the removal of both *nhomie* and *homie* (Fig. 6).

**Figure 6.**
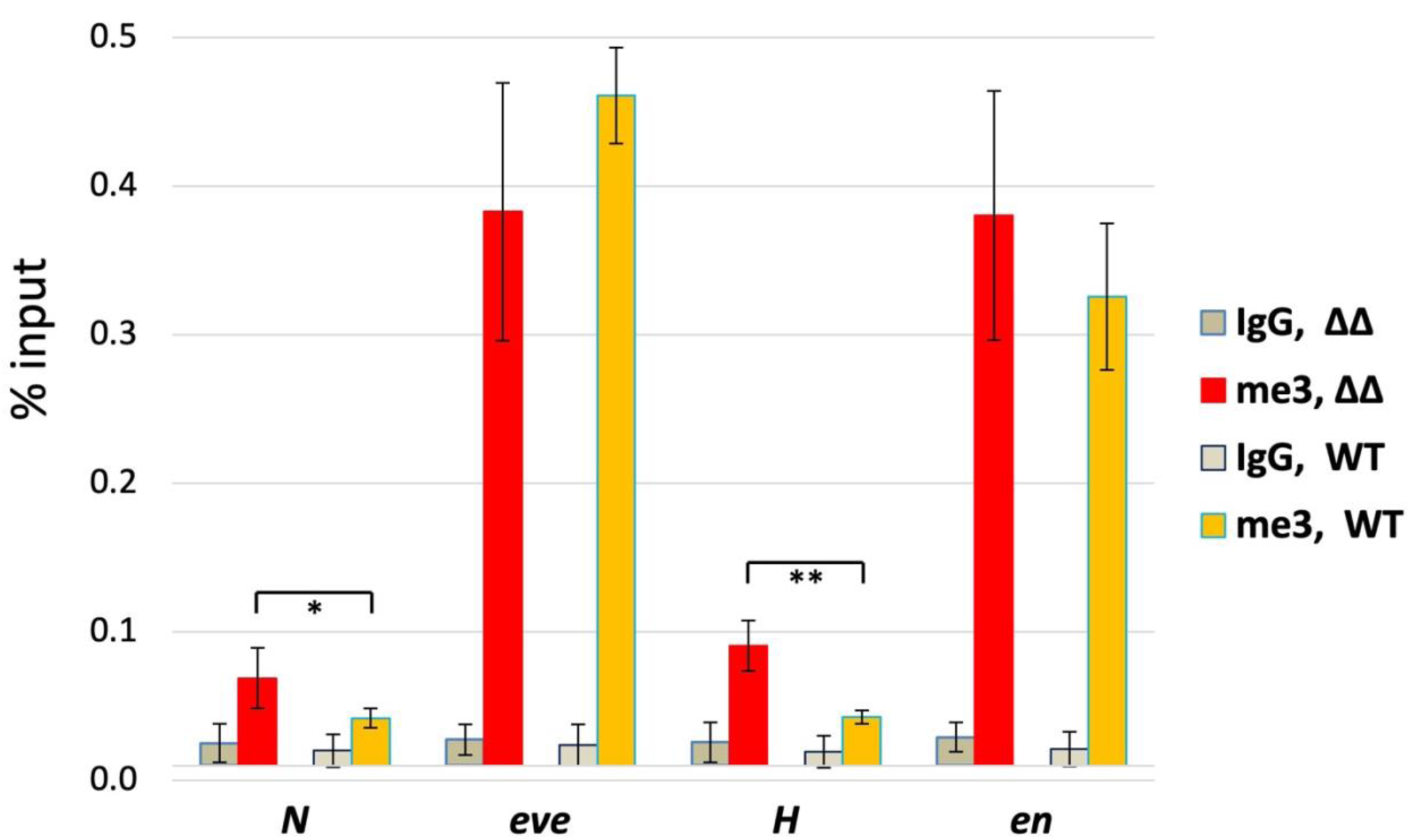
H3K27me3 levels upstream and downstream of *eve* increase when *nhomie* and *homie* are removed. Both Sco/Cy (WT) and ΔNΔH/Cy (ΔΔ) lines were subjected to ChIP analysis using anti-H3K27me3 (me3) antibody or negative control IgG (IgG). Immunoprecipitated DNA were analyzed by qPCR at 4 positions, as follows. N: upstream of the *nhomie* deletion (replaced by λ DNA), *eve*: *eve* coding region, *H*: downstream of the *homie* deletion, *en*: the *engrailed* coding region (positive control). The average and standard deviation from 3 independent experiments are shown as % input. The pair-wise significance of differences (p-values) between anti-H3K27me3 and IgG is indicated as: *: p < .05, **: p < .01.

## DISCUSSION

These studies confirm our model based on transgenes interacting with the intact *eve* locus, in which the transgenic insulator forms a complex with either endogenous *homie* or *nhomie* (or both at the same time). This complex is directional, so that sequences flanking the insulator complex can be either induced to interact or not, depending on whether they are on the same side of the complex in 3-D space. This model requires that the interacting insulators not cause the underlying DNA sequences to be distorted far from their relaxed linear configuration, so that those sequences that end up on opposite sides of the complex are those that are nearest the opposite ends of each insulator. This can be contrasted with how DNA bending protein complexes cause the DNA to fold back on itself to form an enhanceosome [47, 48], or how the TATA box-binding protein complex induces a “kink” in the double helix to facilitate RNA Polymerase II entry [49, 50], or how a nucleosome core particle “wraps” the DNA around itself to form a nucleosome [51]. This “linearity” rule seems to apply not just for *homie* and *nhomie*, but also for other “natural” insulators such as those from the *Abd-B* region of the BX-C, and for multimers of individual insulator binding proteins in studies involving insulator bypass and insulator competition assays [20–22]. The models used to explain the results in those cases presuppose that the interacting DNA sequences maintain a more-or-less linear configuration, so that sequences at the two ends of each insulator end up on opposite sides of the complex, effectively separating them, and those sequences within a few hundred base pairs of the ends, from each other, as well as separating them from those sequences near the “opposite end” of a pairing partner insulator.

In this study, we showed that endogenous *homie* and *nhomie* are necessary for long-range pairing between the *eve* locus and transgenes placed 142kb upstream (Fig. 2). We also showed that long-range trans pairing by transgenic copies of either *homie* or *nhomie* occurs when *homie* and *nhomie* are both absent in cis, but present in trans (Fig. 2). These studies also more directly test whether interacting insulators do so in competition with other possible interactions along a chromosome. When a transgenic insulator has a choice of either of the two endogenous *eve* insulators as a partner, the results are distinctly different than when only one endogenous *eve* insulator is present in the chromosome (Fig. 3). In the latter case, some enhancer-promoter interactions are weakened, consistent with the loss of an interaction with one end of the *eve* locus, while others are strengthened: specifically, those that involve enhancer elements that are closer to the remaining insulator. This strengthening suggests that the remaining insulator may form a more stable or a more frequently forming complex when the other endogenous insulator is missing. This, in turn, is suggestive of a competition occurring between the two endogenous insulators for the transgenic copy when both endogenous insulators are present. This model is consistent with recent genome-wide studies of Drosophila insulator interactions [52], which suggest that dynamic, pairwise insulator interactions are much more common than stable, multi-insulator associations.

There is another likely explanation or contributing factor for the observed changes in *eve* enhancer interactions with a transgenic promoter when one endogenous insulator is missing. If transgenic and endogenous promoters are competing for enhancer activity, then anything that reduces endogenous promoter access is expected to increase transgenic reporter gene (in this case, *lacZ*) expression. We showed here that removing either *homie* or *nhomie* from the *eve* locus reduces *eve* function at early stages, and this likely occurs through a small decrease in expression in early stripes. This may allow enhancers to access the *lacZ* promoter more efficiently and increase reporter gene expression. Additionally, the *eve* locus no longer forms the endogenous *homie-nhomie* interaction loop. This might reduce endogenous *eve* promoter access to the more distant enhancers, as they are further away, on average, than they are when the two endogenous insulators are paired. This, in turn, might reduce competition between the *eve* and transgene promoters, allowing greater reporter gene expression.

There are also other possible factors contributing to the increase in transgene interactions with a single remaining insulator. For example, Micro-C data showed that the endogenous *homie* region interacts with the endogenous promoter, as well as with the *nhomie* region [23], and disruption of these interactions by deleting either *homie* or *nhomie* may well reduce the efficiency of endogenous E-P interactions, allowing greater expression of the transgenic promoter. Another possible model is that a 3-way complex of the transgene with both endogenous insulators might be a larger, and therefore more disruptive, complex than when only a pairwise interaction is possible (when one endogenous insulator is missing), and this could introduce constraints on enhancer-transgene promoter interactions that are relieved when the insulator complex involves only two partners (one endogenous and one transgenic insulator).

We observed that transgenic reporter expression is stronger in late-stage embryos when a trans copy of the *eve* locus is absent (due to the chromosomal deficiency *BSC158*). However, when the *ΔNΔH* chromosome was in trans, the expression was not increased (Fig. 4). Therefore, the effect is probably not due to “trans competition” between the cis copy of the endogenous locus and either the transgene or the trans copy of the endogenous insulators. It is likely due instead to the lack of homologous chromosome pairing when the large *BSC158* deficiency is present in trans. This may cause an increased chromosome “flexibility” at later stages when the homologs are unpaired over the region with the two interacting insulators in cis, leading to a more efficient long-range cis interaction [53]. The *BSC158* deficiency includes about 2/3 of the intervening region between *eve* and the transgene [34, 35]. The reason that the presence of a trans copy of the endogenous locus might not detectably reduce pairing with the transgene could be that the trans interaction is too infrequent or too unstable to reduce cis pairing significantly. So, pairwise interactions may be considerably more stable, or more frequently forming, compared to 3-way or 4-way interactions.

The current studies also establish the function of endogenous insulators in facilitating the full levels of expression of the “insulated” transcription unit. For the *eve* locus, this activity manifests itself as an increase in embryonic defects that are characteristic of reduced *eve* function at a critical early stage (Figs. 5, S5, and S6), when pattern formation is highly dependent on properly formed “pair-rule” *eve* stripes [27, 37–40, 54]. The establishment of fully functional parasegment boundary spacing is known to be highly sensitive to the relative levels of *eve* [41, 42] and its “complementary” pair-rule partner gene *fushi tarazu* (*ftz*) [40, 55]. Specifically, the positioning of the 14 stripes of the segment polarity gene *engrailed* (*en*) become abnormally spaced when either *eve* function is reduced [37–41] or *ftz* function is increased [56], with each odd-numbered *en* stripe becoming closer to the next (posteriorly adjacent) even-numbered *en* stripe. This effect occurs in heterozygotes for *eve* null mutations, yet this abnormal spacing is almost always corrected as pattern formation continues, and few abnormal embryos are seen at the end of embryogenesis (Figs. 5, S5, and S6). However, when *eve* function is reduced further, a critical point is reached where the narrow spacing is no longer efficiently corrected, and defects increase significantly. This is seen as “skips” in the pattern of denticles within even-numbered abdominal segments (this is the phenotype for which *eve* is named, which was based on the initially isolated, strong hypomorphic allele) [27, 54]. We conclude from these results that *homie* deletion alone causes segmentation defects in a sensitized background with one copy of endogenous *eve* function. Deletion of *nhomie* in addition to *homie* increases the severity of the phenotype, indicating that proper activation of *eve*, or efficient interaction between enhancers and the *eve* promoter, requires both *homie* and *nhomie*. We note that because our *homie* deletion is imprecise, the extra deleted sequence could contribute to its more severe phenotype relative to deleting *nhomie* alone. For example, the deletion of the *eve* 3’ PRE might reduce the stability of *eve* enhancer-promoter interactions. Nonetheless, our results indicate that the *eve* insulators, both together and individually, do contribute to the level of expression and function of the *eve* locus.

Finally, we have found that the endogenous *eve* insulators are necessary to “contain” repressive chromatin within the *eve* domain, presumably thereby facilitating the full levels of expression of neighboring transcription units. We previously showed that *homie* has this function in the context of an *eve* “pseudo-locus” transgene [46]. We now show that there is a similar spreading of the *eve* Polycomb domain in both directions in the absence of the endogenous *eve* insulators, measured as an increase in the characteristic histone modification written by Polycomb Repressive Complex 2 (Fig. 6). While these two functions of the *eve* insulators, facilitating full *eve* function and preventing the spread of repressive chromatin, produce incompletely penetrant defects on their own, they are likely to be a sufficient driving force for the evolution of this pair of specifically and strongly interacting insulators. Similarly subtle yet important functions of insulators are likely the rule genome-wide.

## Materials and Methods

### Creation of *nhomie* and *homie* deletions, and breakpoint analysis of chromosomal deficiencies

Creation of the *ΔN* line has been described previously [23]. In short, we inserted two attP sites flanking endogenous *nhomie*. To do this, we used a donor plasmid, P-attPx2-*mw,* carrying two 102bp attP sequences flanking a modified *mini-white* (*mw*) gene. The following modification was made to *mw*: the Wari insulator [57] was deleted from the standard *mw,* and Glass binding sites were added to boost the eye color. The endogenous *nhomie* region was replaced with this attP-*mw*-attP using CRISPR. This chromosomal modification resulted in one attP site being inserted in the intron of *CG12134*, and the other being inserted between the *eve* 7-stripe enhancer and the 3+7 stripe enhancer (Fig. 1C). This process also deleted 2.2kb of endogenous sequence, including *nhomie* and the *eve* 7-stripe enhancer. After identifying a successful insertion (*NattPmw*), *mw* was replaced by the same 2.2kb sequence, but with 600bp of phage λ DNA in place of 600bp *nhomie*, using recombinase-mediated cassette exchange (RMCE) [58].

Several attempts to insert attP sites flanking *homie* by CRISPR using different gRNA sets failed. While we do not fully understand why our multiple trials of this strategy did not work, it is possible that removing *homie* is either dominantly lethal, or that it reduces viability such that recovery of the resulting chromosome is difficult, perhaps due to the function of *homie* in preventing repression of the adjacent, essential gene, *TER94*. However, we have a homed transgenic line that inserted at +9111bp relative to the *eve* transcription start site [24]. Using this line, we successfully mobilized the transgene, creating a small deletion (2R:9987986– 2R:9989354 fusion, the two ends share the sequence AAA) (Fig. 1B, sequence is shown in Fig. S1). The deletion includes the *eve* 3’ PRE, *homie*, and the first exon of *TER94* (Fig. 1B). We described this line as *ΔH*. *ΔH* is homozygous lethal. The same strategy that was used to make the *nhomie* deletion on a wt chromosome was used to modify *nhomie* on the *ΔH* chromosome, creating the *ΔNΔH* deficiency chromosome (Fig. 1C). As expected, *ΔNΔH* is homozygous lethal. Lines carrying either the *ΔH* or *ΔNΔH* chromosome over a balancer were self-crossed to make *ΔH* and *ΔNΔH* (“*ΔΔ*” in some figure labels) homozygous embryos, respectively.

For *Df(2R)eve* [27, 36], information in Flybase [34] suggested that its breakpoints are in *Mef-2* and *TER94*, so we tested potential PCR primers to find ones that amplified junction fragments. These PCR products were then sequenced to identify the specific breakpoints (Fig. S1). Primers used for PCR fragments that were used for sequencing analysis are following: For *ΔH* deletion, CAGTCGAGCCTCCGTAAGGG and CCTCCAGCAAAGGATGACTTG. For *Df(2R)eve*, TTTCAACCGCACACAATCC and CATTCATTCCAAATCACGCAC.

### Creation of attP sites at –142kb and insertion of reporter transgenes

Two attP sites were inserted near the original −142kb attP site [16, 24] on the wt, *ΔN*, *ΔH*, and *ΔNΔH* chromosomes, using the same CRISPR strategy described above. Then, *mini-white* was replaced with each of the reporter transgenes using RMCE. Dual reporter transgenes were used (Fig. 1D) [16], carrying either 400bp *homie* (*Z-homie-G*), 600bp *nhomie* (*G-nhomie-Z*), or 500bp of *lambda* DNA (*Z-lambda-G*). The orientation in which the *lacZ* reporter is more strongly activated due to orientation-specific long-range pairing was used in this study.

### Analysis of transgenic lines

*In situ* hybridization was performed based on previously published methods [59], except that RNA was visualized using a histochemical reaction, as described previously [13]. Briefly, approximately 60μl embryos per sample (stage 4-15 embryos are present in each sample) are subjected to the process. DIG-labeled antisense RNA probes against *lacZ* or *GFP* were visualized using alkaline phosphatase-conjugated anti-DIG antibody (Roche), using CBIP and NBT as substrates (Roche). Once color was developed based on positive control expression, the reactions of all samples were stopped simultaneously. Each set of experiments was carried out with the positive control and experimental samples in parallel to minimize experimental variation. Each experiment was performed more than twice, with independent *in situ* procedures. Representative expression is shown in the figures.

### Analysis of embryonic cuticle defects

To identify segmentation defects in developing embryos, embryos were collected and analyzed as described previously [23], with minor modifications. Briefly, embryos were collected for 2.5-3 hrs, allowed to develop for 20–21 hr at 25°C, then dechorionated and mounted in a 1:1 mixture of Hoyer’s medium and lactic acid. Mounted embryos were left at room temperature (RT) until they cleared, and the patterns of ventral abdominal denticles were examined and tallied as follows. Loss of at least one-fifth of a denticle band in A1-A8 was counted as ‘missing’. Fused denticle bands, which rarely occurred, were also counted as a ‘missing’ band. Minor defects such as those within individual denticle rows were not counted.

Wild-type (wt), *ΔN, Sco/SM6a,Cy, ΔH/SM6a,Cy*, and *ΔNΔH/SM6a,Cy* lines were self-crossed for Fig. S4. For each cross, 3-4 independent cuticle preparations were analyzed. To sensitize for functional differences of these chromosomes, each of the lines was crossed with each of the *eve* heterozygous mutant lines *Df(2R)BSC158/CyO* (Figs. 5, S6A, and S7A), *Df(2R)eve/CyO* (Figs. S6B and S7B), and *eve^R13^/CyO* (Fig. S6C, S7C). The two directions of each cross were analyzed separately. For each cross, 5-8 independent cuticle preparations were analyzed.

In the sensitized condition, segmentation defects were quantified in two ways: the percentage of embryos with segmentation defects (% affected) and the average number of ventral abdominal denticle bands deleted per non-*wt* embryo (severity score). The data from each set of cuticle preparations are presented as box-and-whiskers plots (Figs. 5, S6), and the pair-wise significance of differences (p-values) between lines was calculated using the t-test function (2-tailed, unequal variance) in Excel (Microsoft). For Fig. 5, two sets of data were combined, one directly comparing WT, *ΔN*, and *ΔH* (shown in Fig. S6A) and the other directly comparing WT, *ΔH*, and *ΔNΔH*. The raw data from this 2nd comparison were used in Fig. 5, along with scaled data for *ΔN* from the 1st comparison. For this purpose, individual data points from the 1st comparison were scaled based on their values relative to WT and *ΔH* (they were all between the average values for WT and *ΔH* in the 1st comparison, and were scaled to have the same fractional distance from the average WT and *ΔH* values in the 2nd comparison as they did in the first comparison). The statistical analysis for Fig. 5 used these scaled values for *ΔN* in combination with the raw data for the other lines. For a different visual comparison, stacked graphs are shown in Fig. S7. For this, the percentages of embryos (average and standard deviation) with 4 different severities of phenotype are shown: wild-type, mild (1 band missing), moderate (2 bands missing), and severe (3-4 bands missing).

### Analysis of H3K27me3 levels

ChIP-qPCR was performed based on previously published methods [60]. Briefly, stage 4-17 embryos (approximately 200ul) from *Sco/Cy* and *ΔNΔH/Cy* were cross-linked in 2% formaldehyde for 10 min. After sonication, 50μg of isolated chromatin was used to immunoprecipitate with anti-H3K27me3 (Millipore), with rabbit IgG (Jackson ImmunoResearch) used as the negative control. Precipitated chromatin samples were collected using ProteinG magnetic beads. After reversing cross-links, purified samples were dissolved in 50 μl TE, and 1 μl was used for each qPCR reaction. Triplicate samples were analyzed by real-time PCR (Life Technologies, StepOne Plus), using SYBR Green Master Mix with ROX dye (Roche Applied Science). Data were analyzed with StepOne software (Life Technologies), using the standard curve method. The data are presented as average % inputs from 3 independent ChIP analyses. Standard deviations were calculated using Excel software (Microsoft). The pair-wise significance of differences (p-values) between lines was calculated using the t-test function in Excel (Microsoft).

Primers used for qPCR were as follows. *N* (upstream of *nhomie*) : CGGAGAATCCGGCATTGTTA and GCTTGCGTGATTTCTTCTCC, *eve* (*eve* coding region): TCCAGTCCGGATAACTCCTTG AAC and TGTAGAACTCCTTCTCCAAGCGAC, *H* (downstream of *homie*): AAGGGCCACATCGCAGACATACTA and GTCGCGGTAAATGTCTTTGTCTCG, *en* (*engrailed* coding region): GAGAACCAGGCCAGCATATT and CTAAACTCCAGCAGATCCACTC.

## Acknowledgements

We thank Qing Liu and Lanxi Li for excellent technical assistance. We also thank Flybase and the Bloomington Drosophila Stock Center, which were instrumental in this work.

## Supplemental Figures

**Figure S1.**
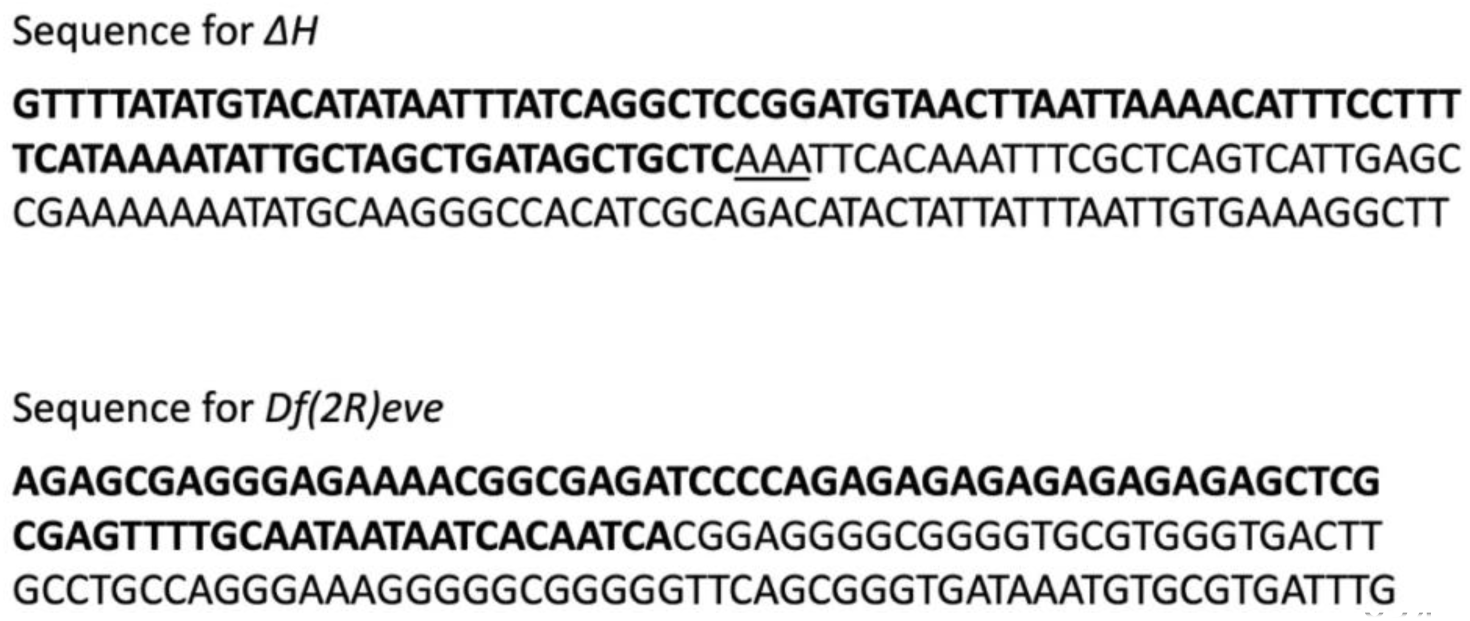
Junction sequences of the chromosomal deletions *ΔH* and *Df(2R)eve*. Sequences upsteam of each junction are shown in boldface. The underlined AAA sequence in *ΔH* could be from either side of the junction, since it is found on both sides.

**Figure S2.**
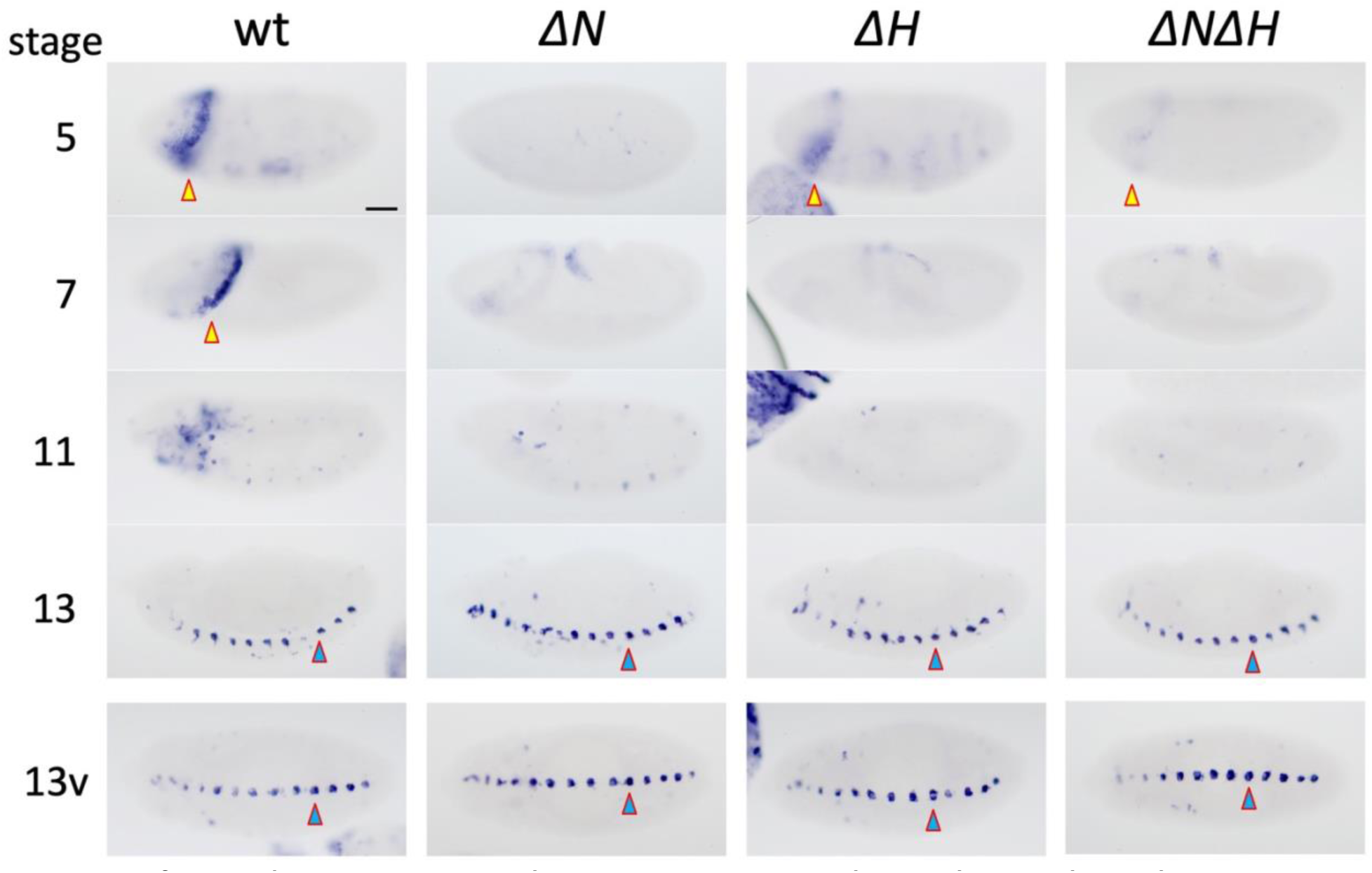
λ DNA does not promote long-range pairing, either with or without the endogenous *eve* insulators. *lacZ* expression from the reporter gene *Z-lambda-G* on wt, *ΔN*, *ΔH*, and *ΔNΔH* chromosomes. Note that there is no *eve*-like expression. Consistent with previous studies [13, 16], non-*eve* like expression is seen in the form of head stripes (yellow with red outline) and *hebe*-like ventral mid-line expression (blue with red outline). Embryonic stages 5, 7, 11, and 13 are shown (ventral views in bottom row). Scale Bar: 50μm.

**Figure S3.**
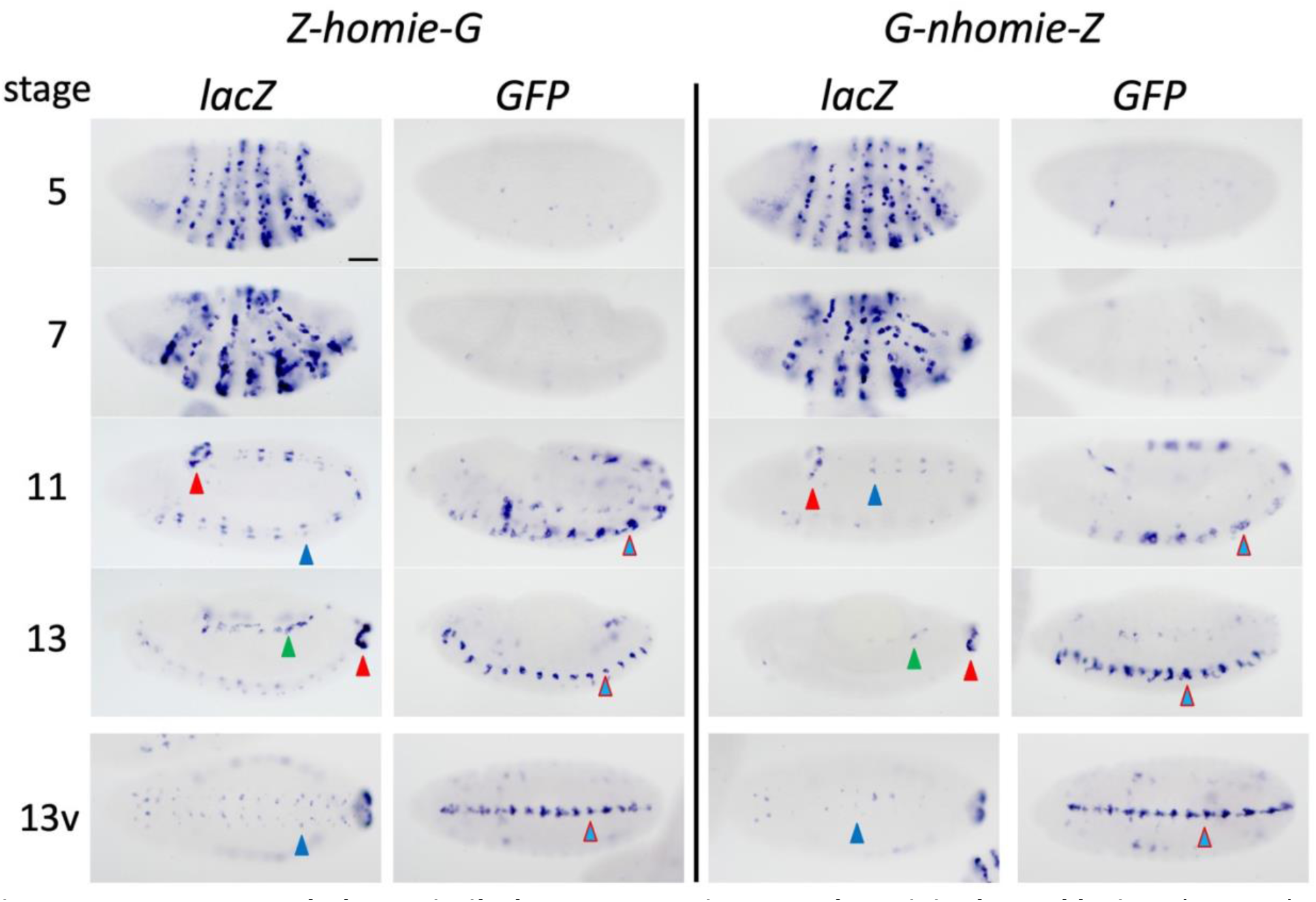
Transgenes behave similarly at an attP site near the original –142kb site. *lacZ* and *GFP* expression from *Z-homie-G* and *G-nhomie-Z*. Consistent with a previous study [16, 25], LR pairing is biased toward one transgenic reporter or the other, depending on the orientation of *homie* or *nhomie* in the transgene (see main text). Embryonic stages 5, 7, 11, and 13 are shown (ventral views in bottom row). Scale Bar: 50μm.

**Figure S4.**
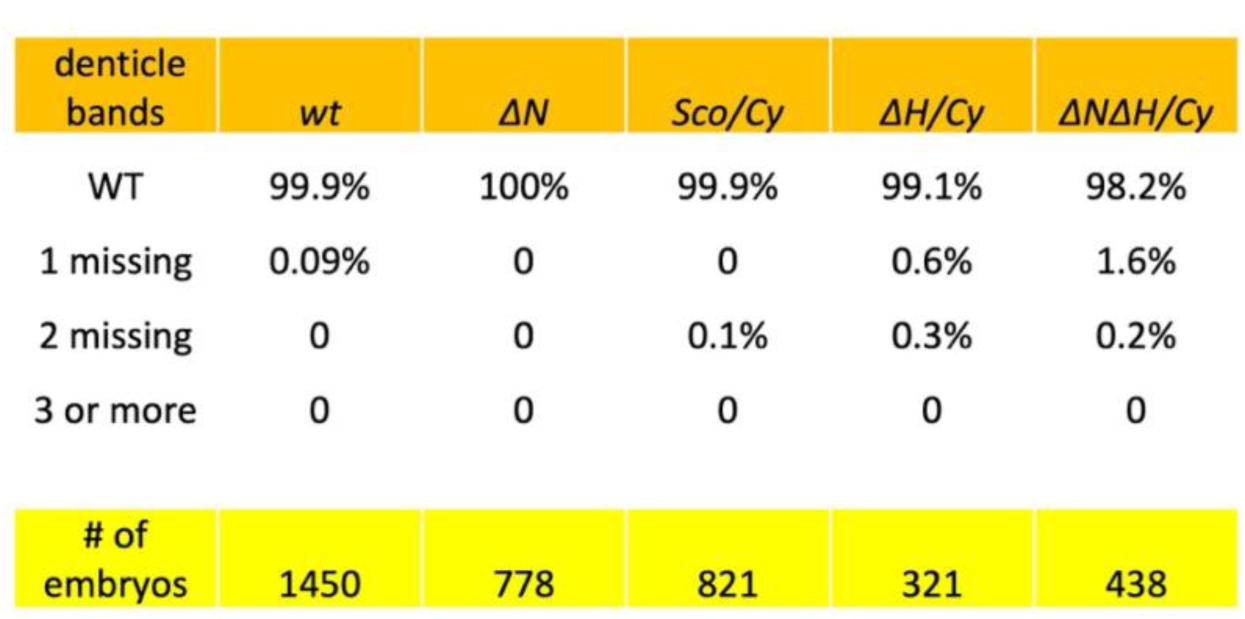
Embryos homozygous for the *homie* deletion and for the *homie*+*nhomie* deletion show few cuticle defects. Lines of *wt, ΔN, Sco/Cy, ΔH/Cy,* and *ΔNΔH/Cy* were self-crossed, and cuticle defects were counted as either WT (no missing denticle bands), or as having 1, 2, or 3 or more missing denticle bands. For *ΔH* and *ΔNΔH*, homozygotes are expected to be 25% of the population. Total numbers of counted embryos are shown at the bottom. Number of cuticle preparations included: wt and *ΔN*: n=4; *Sco/Cy*, *ΔH/Cy*, and *ΔNΔH/Cy*: n=3.

**Figure S5.**
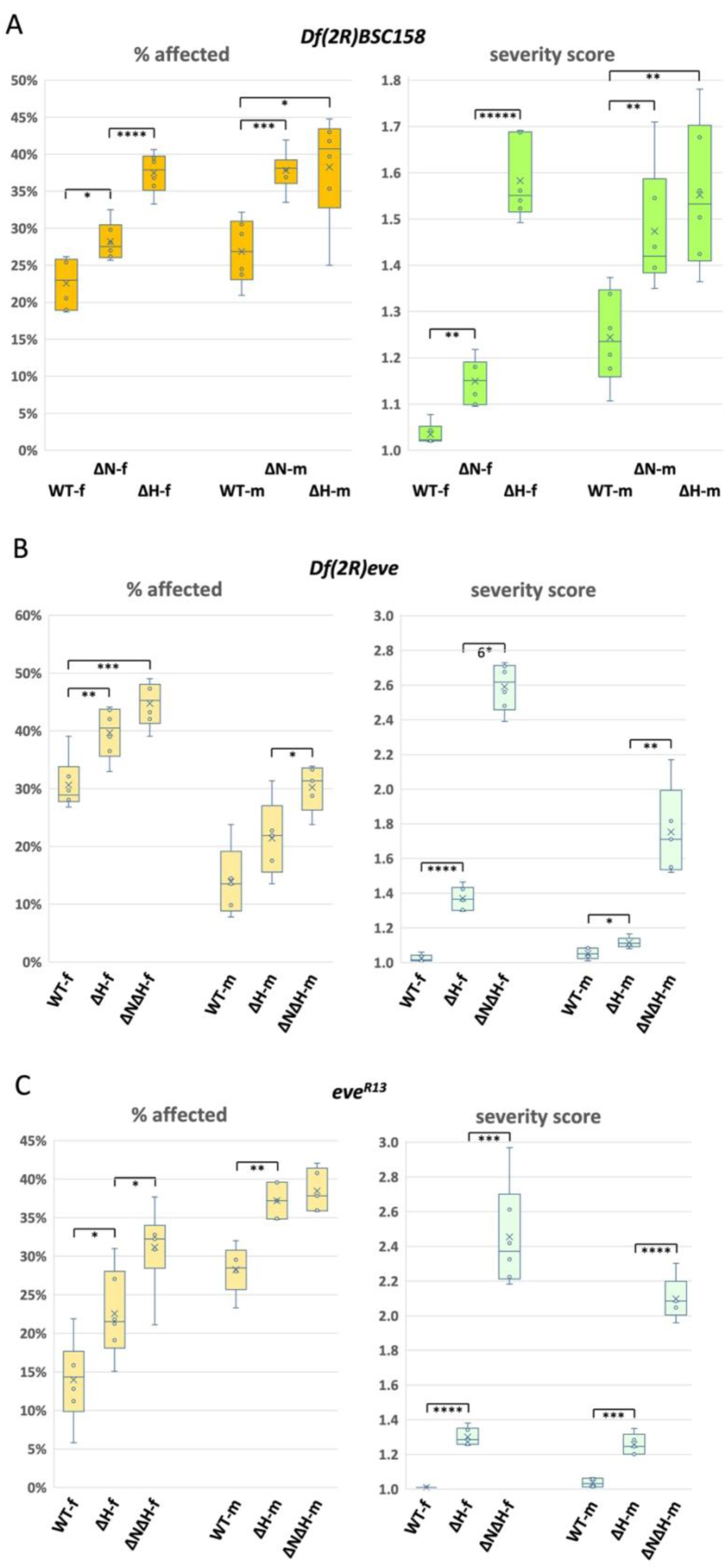
Deleting either *nhomie* or *homie* from the endogenous *eve* locus compromises embryonic function. Embryonic cuticle defects were tabulated from the following crosses: **A.** *Sco/Cy* (WT)*, ΔN/Cy* (ΔN), and *ΔH/Cy* (ΔH) crossed with *Df(2R)BSC158/Cy*. **B.** *Sco/Cy* (WT)*, ΔH/Cy* (ΔH), and *ΔNΔH/Cy* (ΔNΔH) crossed with *Df(2R)eve/Cy.* **C.** *Sco/Cy* (WT)*, ΔH/Cy* (ΔH), and *ΔNΔH/Cy* (ΔNΔH) crossed with *eve^R13^/Cy.* The direction of each cross is indicated as -f: female and -m: male after the *eve*-locus genotype. The percentage of embryos showing deleted ventral abdominal denticle bands (left graph, % affected) and the average number of denticle bands deleted per non-*wt* embryo (right graph, severity score) are each shown as a box-and-whiskers plot. The pair-wise significance of differences (p-values) are shown by the number of asterisks. *: p < 0.05, **: p < 0.01, ***: p < 0.001, ****: p < 0.0001, *****: p < 10^−5^, *^6^: p < 10^−6^. The number of cuticle preparations included (n) and total number of embryos counted were as follows: in **A**, WT-f (n=6, 2722 embryos), ΔN-f (n=6, 1676), ΔH-f (n=6, 3661), WT-m (n=6, 1119), ΔN-m (n=6, 1623), ΔH-m (n=6, 1000); in **B**, WT-f (n=6, 2457), ΔH-f (n=6, 3773), ΔNΔH-f (n=6, 2624), WT-m (n=5, 2123), ΔH-m (n=5, 2320), ΔNΔH-m (n=5, 2217); in **C**, WT-f (n=6, 1457), ΔH-f (n=6, 3434), ΔNΔH-f (n=6, 1500), WT-m (n=5, 2641), ΔH-m (n=5, 2939), ΔNΔH-m (n=5, 3289).

**Figure S6.**
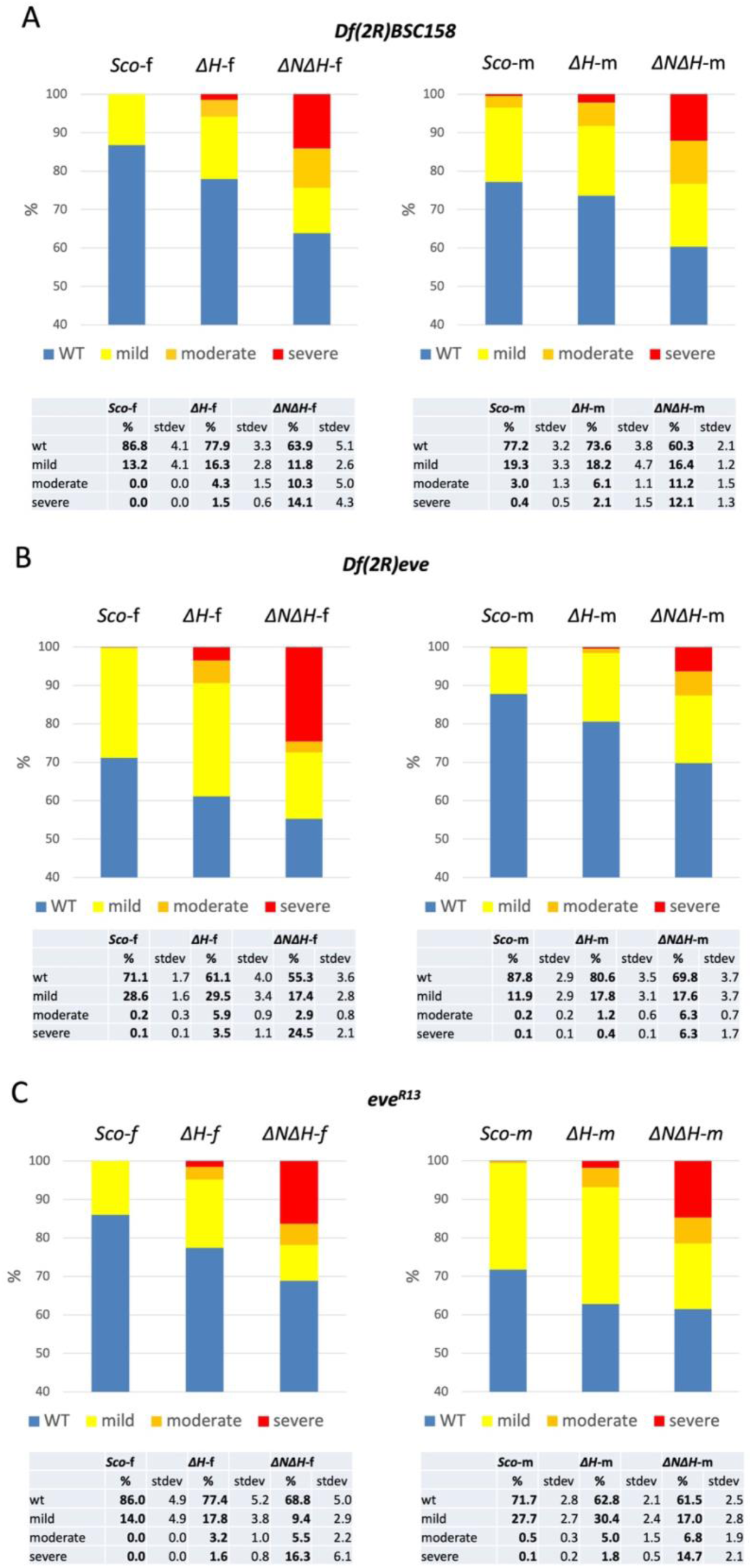
Deleting both *nhomie* and *homie* from endogenous *eve* compromises function more than does deleting only *homie.* *Sco* (wt control from *Sco/Cy* stock)*, ΔH/Cy* (*ΔH*), and *ΔNΔH/Cy* (*ΔNΔH*) were crossed with either, in **A**, *Df(2R)BSC158/Cy*, or in **B**, *Df(2R)eve/Cy*, or in **C**, *eve^R13^/Cy.* The same data used for Figs. 5, S5B, and S5C are shown here as stacked graphs. WT: no missing ventral denticle bands; mild: 1 missing; moderate: 2 missing; severe: 3-4 missing. Tables at the bottom show the average percentages of embryos in each deficiency class (%) and their standard deviations (stdev).

**Figure S7.**
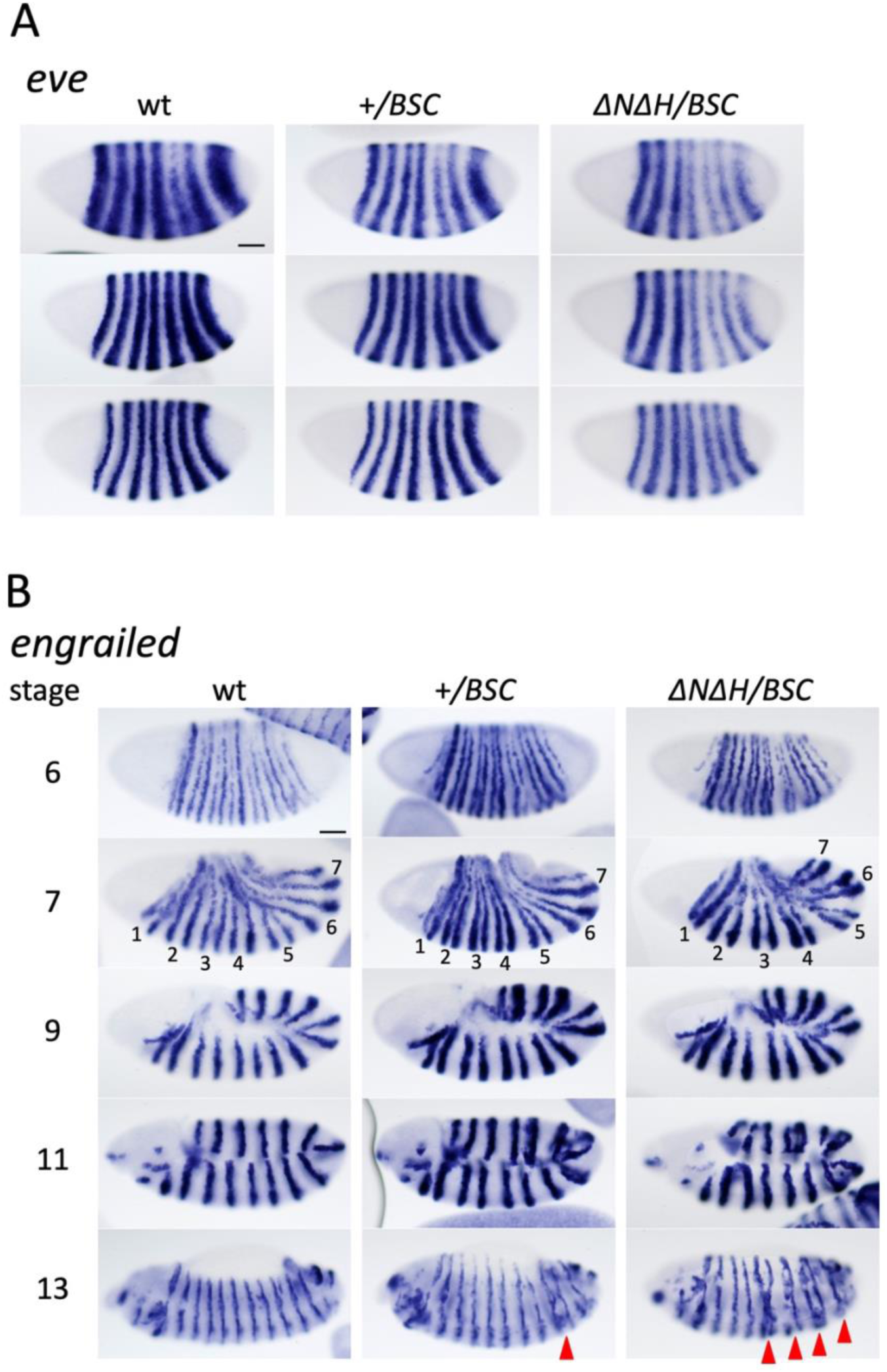
Abnormal *eve* and *engrailed* expression is seen in *ΔNΔH/Df(2R)BSC158* embryos. **A.** *eve* expression at stage 5 (early to later stages are shown from top to bottom). **B.** *engrailed* expression at stages 6, 7, 9, 11, and 13. At stage 7, the positions of early *eve* stripes are numbered. Fused *engrailed* stripes that likely prefigure missing denticle bands are marked with red arrowheads at stage 13. wt: homozygous wildtype genotype (both copies of the *eve* locus are intact), +*/BSC*: wt over *Df(2R)BSC158* (1 copy of the *eve* locus present), *ΔNΔH/BSC*: *ΔNΔH* over *Df(2R)BSC158* (1 copy of the *ΔNΔH eve* locus present). Scale Bar: 50μm.

